# Spatial transcriptomics exploration of the primary neuroblastoma microenvironment unveils novel paracrine interactions

**DOI:** 10.1101/2024.12.21.629891

**Authors:** Joachim T. Siaw, Peter Merseburger, Marcus Borenäs, Caroline Jansson, Jenny Karlson, Arne Claeys, Eva Jennische, Dan E. Lind, David Gisselsson Nord, Ruth H. Palmer, Jimmy Van den Eynden

## Abstract

High-risk neuroblastomas exhibit a high degree of intratumoral heterogeneity. Single-cell RNA sequencing has greatly improved our understanding of these tumors, but the method lacks cellular tissue context and spatial information about local signaling dynamics. To address this gap, we profiled untreated and chemotherapy-treated high-risk neuroblastomas from 2 patients using spatial transcriptomics. We confirmed the transcriptional and cellular heterogeneous nature of the neuroblastoma microenvironment and identified several unique spatial niches and patterns. In one of the treated tumors, a spatially constrained cluster of undifferentiated and 11p-gained cancer cells was identified, surrounded by a rim of macrophages. A signaling interaction between the chemokine *CCL18* and its receptor *PITPNM3* was predicted between these cells and we experimentally demonstrated that *CCL18* increases neuroblastoma cell migration. In the other tumor, we identified a stromal cluster with high transcriptional similarity to the adrenal cortex. These adrenocortical-like cells expressed the ALK ligand *ALKAL2* and were predicted to communicate with neighboring *ALK* expressing cancer cells. We demonstrated a unique developmental pattern of adrenal medulla-specific expression of *ALK* and adrenocortical-specific expression of *ALKAL2*, suggesting a role of these signaling interactions in neuroblastoma carcinogenesis.

## INTRODUCTION

The pediatric cancer neuroblastoma (NB) exhibits both clinical and biological diversity, with distinct clinical outcomes that range from spontaneous remission to relentless disease progression^1,2^. High-risk NB is characterized by a high degree of inter- and intratumoral heterogeneity (ITH) which are considered major and independent contributors to disease progression and treatment failure^3^. A key contributor to intertumoral heterogeneity is the tumor’s site of origin. Tumors originating in the adrenal gland are the most common and tend to be more aggressive compared to those from non-adrenal sites, such as the thorax^4^. Earlier efforts to untangle ITH primarily relied on multi-regional tumor sampling and whole genome (WGS) and exome sequencing (WES)^3,5–7^.

More recently, single cell (sc) RNA-Seq has vastly enhanced our understanding of NB ITH. It has spearheaded the discovery of new cell types and cell states, reflecting the underlying heterogeneity and plasticity within tumor tissues^8–11^. NB is thought to originate from neural crest-derived precursor cells and cellular intermediates along the sympathoadrenal developmental trajectory. These cancer cells exhibit a continuum of identities, spanning the transcriptional space from early neural crest-derived precursors to immature and mature sympathoadrenal lineage-committed cells^2,4,9,10,12,13^. The tumor cells are believed to exist between two interconvertible transcriptional cell states, namely adrenergic and mesenchymal states, which exhibit different responses to therapy^14,15^. These plastic cell states increase ITH further^5,16,17^. A key methodological step of all sc-RNA studies involves tissue dissociation into single cells, resulting in the loss of spatial information and, in some cases, selective depletion of certain cell populations^8^. Consequently, information on the complex intratumoral microhabitats, their interactions within the tissue and the precise cellular tumor composition is lost.

To better understand the cellular composition of the NB tumor microenvironment (TME) and the specific spatial interactions between these previously identified cell types and states, we spatially profiled the whole transcriptome of 6 sections from 3 NB tumors using the *Visium* gene expression platform. We confirmed the strong cellular and transcriptional heterogeneity of the NB TME and identified several spatially organized neuroendocrine cancer cell populations that were characterized by a strongly varying degree of adrenergic or mesenchymal differentiation. A unique spatial niche of undifferentiated and 11p-amplified cancer cells was identified in a chemotherapy-treated tumor sample that was surrounded by a rim of *CCL18* expressing macrophages. In another chemotherapy-treated NB sample, we identified an adrenocortical-like stromal group of cells, exclusively expressing *ALKAL2*, the ligand of the oncogenic receptor *ALK*.

## RESULTS

### Cellular and transcriptional heterogeneity of the NB tumor microenvironment

Tumor sections were obtained from two NB patients (referred to as *NB1* and *NB2*) with known genomic copy number profiles. Both patients received prior chemotherapy (rapid COJEC protocol, with switch to lorlatinib monotherapy for *NB2* due to poor chemotherapy tolerance; tumor samples referred to as *NB1Post* and *NB2Post*) and for *NB1* we also had pretherapy tumor samples available (referred to as *NB1Pre*). *NB1* was a 2-year-old boy with an abdominal tumor that harbored multiple segmental copy number alterations including 11q loss but without evidence of *MYCN* amplification or *ALK* mutation. *NB2* was a 2-year-old girl with an abdominal tumor that was *MYCN* amplified and had both an *ALK* -p.R1275Q mutation and an *HRAS* mutation found by whole genome sequencing at diagnosis (Fig. 1A). Resequencing of the resection specimen (*NB2Post*) after chemotherapy showed retained *MYCN* amplification and *HRAS* mutation but absence of *ALK* mutation, even after deep NGS panel sequencing (Oncomine Focus panel). Two sections were profiled from each tumor using the 10X Genomics *Visium* spatial transcriptomics (ST) platform, resulting in the analyses of 6 sections obtained from 3 different tumors (Fig. 1B).

**Figure 1.**
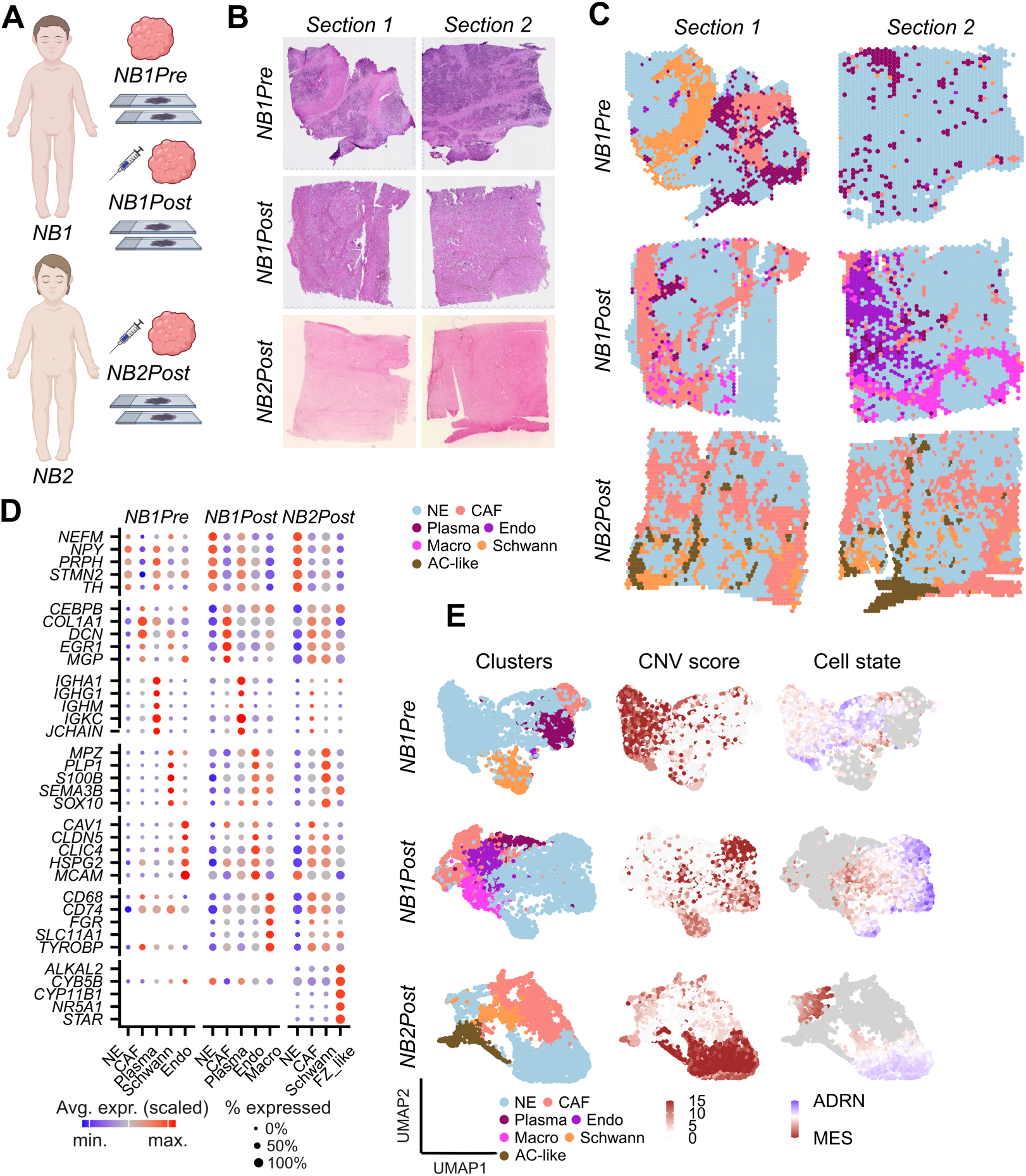
Spatial transcriptomic profiling of 6 tumor sections obtained from 2 NB patients. (**A**) The Visium spatial transcriptomics platform was used to profile 3 tumors (2 sections each) from 2 NB patients (*NB1* and *NB2*). Both patients received prior chemotherapy (*NB1Post* and *NB2Post*) and for *NB1* we also profiled pretherapy tumor materials (*NB1Pre)*. Created in BioRender. (**B**) Hematoxylin and eosin (H&E) staining of the 6 tumor sections that were used in this study. (**C**) Clustering and annotation of 7 main spatial clusters across the 6 samples. Cluster annotations were based on the most representative cell type, as predicted from marker gene expression, enrichment analyses and similarities to single cell data. See Suppl. Fig. 1-2 for details. (**D**) Dot plots showing relative expression (colors) and proportional expression in the spots (sizes) of the top 5 representative genes for each cluster. Genes derived from the leading edges from the GSEA shown in suppl. Fig. 1D. (**E**) UMAP plots showing the main clusters corresponding to each tumor (left), the CNV score, which is representative for the overall copy number variability (middle) and the cell state (adrenergic or mesenchymal as indicated by color key; right). NE, neuroendocrine cells; CAF, cancer associated fibroblasts; Schwann, Schwann cells; Macro, macrophages; Endo, endothelial cells; Plasma, plasma cells; AC-like, adrenocortical-like; ADRN, adrenergic; MES, mesenchymal.

An unsupervised clustering approach resulted in the identification of 11 - 14 distinct spatial clusters in these 3 tumors (Fig. S1). A differential gene expression (DGE) analysis was performed between each cluster and the rest of the sample and clusters were then merged and annotated based on 1) their DGE pattern and the expression of key marker genes (Table S1; Fig. S1C), 2) a gene set enrichment analysis (GSEA) using an extended cell marker gene set derived from PanglaoDB (Fig. S1D), and, 3) their similarity to previously annotated single cell (sc)-RNA data as available in the recently published NBAtlas^18^ (Fig. S2). This annotation resulted in 7 distinct clusters across the 3 tumor samples (Fig. 1C-E).

The first cluster was present in all samples and was characterized by high expression of *TH*, *NPY* and *NEFM*, which are typical neuroendocrine (NE) cancer cell markers (Fig. 1D). Their NE identity was confirmed by the high similarity to scRNA annotated NE cells and a deconvolution analysis predicted NE cells were indeed the dominant cell type (between 51% and 68% for the 3 tumors) in this cluster (Fig. S2A-B). Additionally, a copy number analysis suggested that these cells had the highest copy number variability (i.e., high CNV scores; Fig. 1E), further confirming their NE cancer cell identity. Based on these findings this cluster was annotated as the “NE cluster”. It was present at distinct locations in each tumor and was further subdivided into 2-4 tumor-specific NE subclusters (called NE1-NE4) based on spatial location and expression profile similarities (Fig. S2-3).

Because NE cancer cells are known to occur in 2 differentiation states (mesenchymal and adrenergic), we determined the relative state using a single visualization score based on the log2 ratio of the normalized ADRN and MES UCell scores (see *Methods*). Interestingly, the individual NE spots filled a wide spectrum between mesenchymal and adrenergic cell states in each tumor (Fig. 1E) and, strikingly, *NB2Post* contained one spatially and transcriptionally clearly distinct mesenchymal NE subcluster (NE2, Fig. S4A). This cluster was further characterized by low copy number variability (Fig. 1E), showed poor similarity to single cell NE cells (although *TH* was overexpressed and it was clearly enriched for neuronal cells; Fig. S1C-D, Fig. S3C) and, interestingly, was the only cluster that was enriched for chromaffin (CA) cell markers (Fig. S1D). Additionally, the most upregulated gene was *PNMT* (Table S1; Fig. S3D), which encodes an enzyme that methylates norepinephrine to form epinephrine, a key role of adrenal chromaffin cells^11,19^.

The second cluster was similarly present in all samples and had high expression of several collagen-and extracellular matrix-related related genes (e.g., *DCN*, *COL1A1, MGP;* Fig. 1D). It was most similar to scRNA annotated fibroblasts, which were also the most dominant cells (56% - 73%; Fig. S2A-B). Based on these findings this cluster was annotated as the “cancer-associated fibroblast (CAF) cluster”.

The other 5 clusters were only identified in one or two tumor samples and their similarities to scRNA-derived cell types were less delineated with deconvolution analyses indicating more heterogeneous populations (Fig. S2). These clusters were annotated as “Macro”, “Plasma”, “Schwann” and “Endo” based on the expression of macrophage, plasma cell, Schwann cell and endothelial cell markers, respectively (Fig. 1D).

### Identification of a unique adrenocortical-like group of cells in NB

The last cluster was only present in the *NB2Post* sections. A comparison to the NB scRNA reference resulted in the highest similarity with cells that were previously annotated as adrenocortical zona glomerulosa (ZG) cells (Fig. S2A). When extending the enrichment analysis to the Reactome gene set, strong enrichments were found for genes involved in steroid (hormone) metabolism (Fig. 2A and Fig. S5A), and, indeed, many genes known to be involved in steroidogenesis were almost exclusively expressed by this cluster (e.g., *NRA5A1, CYP11B1*, *STAR* ; Fig. 1D and Fig. 2B-C). A unique spatial organisation of this tissue was observed. On the one hand a spatially distinct cluster was observed at the bottom of both sections of *NB2Post*, adjacent to the more centrally located NE2-CA clusters (Fig. 3A). On the other hand, both samples contained a combination of stripes and nests of steroid gene expressing cells/spots that were scattered throughout the tumor tissue (Fig. 1C and Fig. 2B-C).

**Figure 2.**
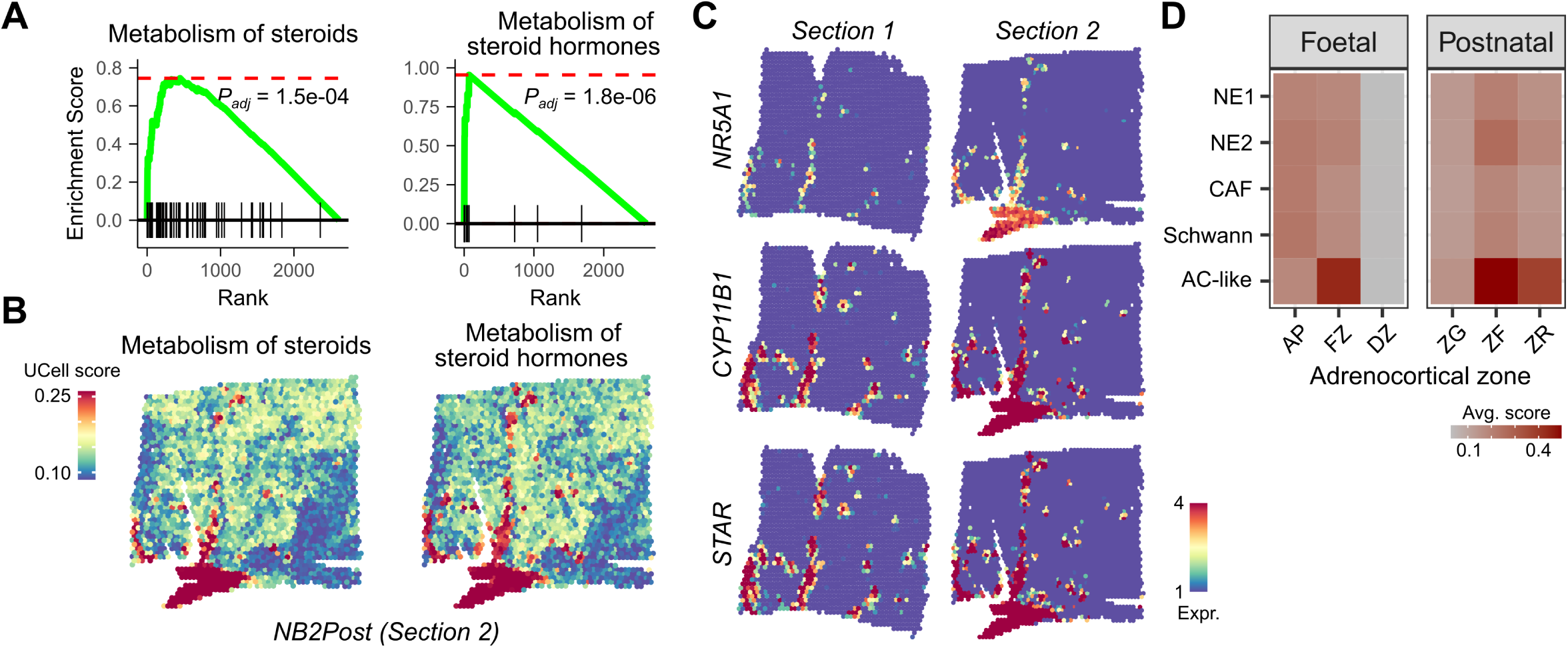
Identification of an adrenocortical-like cell cluster in the NB tumor landscape. Analysis of the cluster annotated as AC-like in *NB2Post* tumors. (**A**) Gene set enrichment running score plots for Reactome gene sets “Metabolism of steroids” and “Metabolism of steroid hormones” as indicated. *P* value calculated using a permutation test as implemented in the *clusterProfiler* R package. See table S2 for complete GSEA results. (**B**) Spatial feature plots showing the expression of these Reactome gene set signatures (UCell scores) in *section 2*. (**C**) Spatial feature plots of 3 key adrenocortical marker genes as indicated. (**D**) Heatmaps showing the average UCell scores of fetal and postnatal adrenocortical cell type signatures on the main clusters identified in *NB2Post*. AP, adrenal primordium; FZ, fetal zone; DZ, definitive zone; ZG, zona glomerulosa; ZF, zona fasciculata; ZR, zona reticularis.

**Figure 3.**
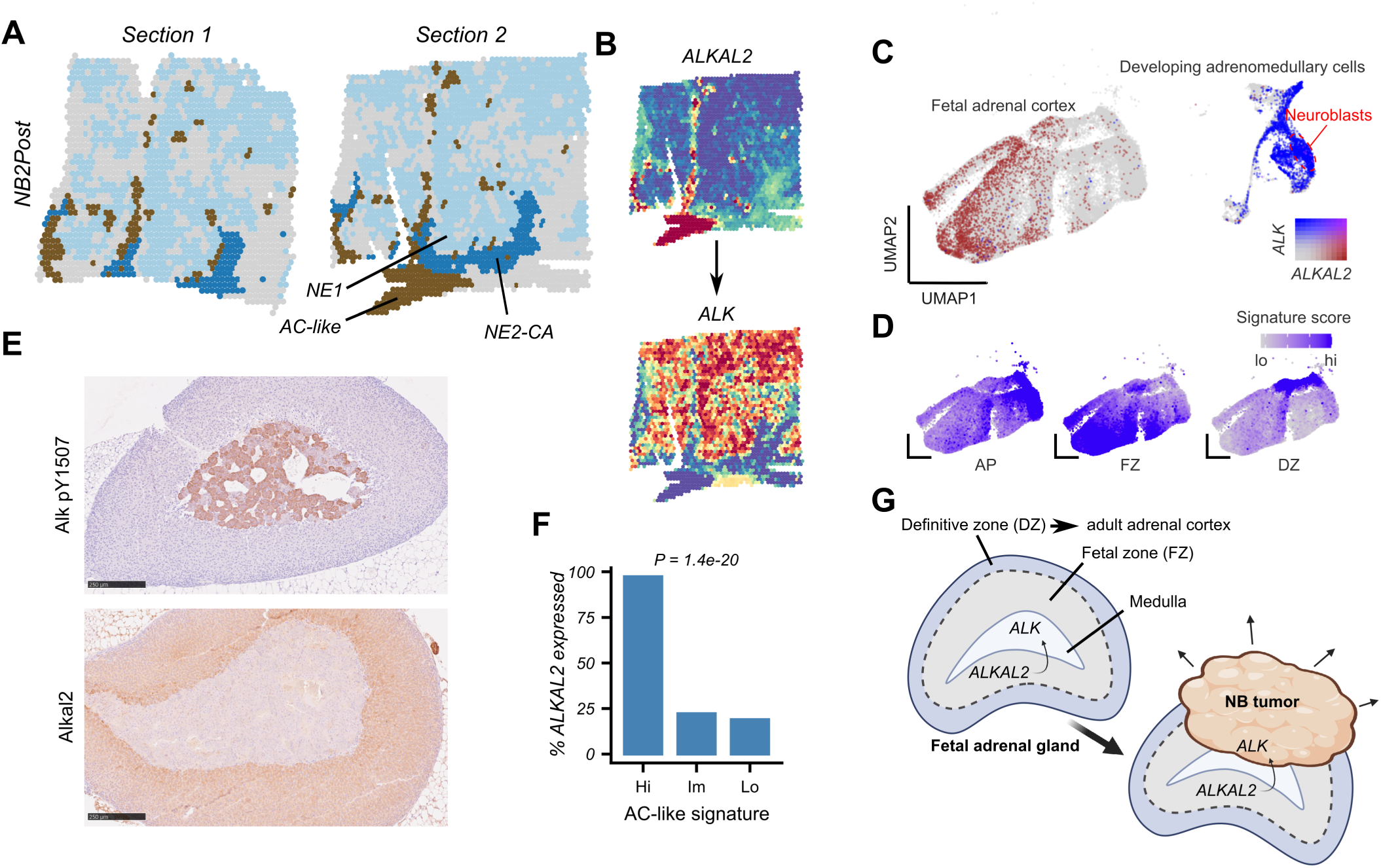
Complementary spatial expression patterns between *ALKAL2* expressing adrenocortical-like cells and *ALK* expressing tumor cells in *NB2Post* tumors. **(A)** Tumor tissue locations of the AC-like cells, NE1 cancer cells and NE2-CA cells in both sections of *NB2Post* tumors as indicated. (**B**) Spatial feature plots showing corresponding *ALKAL2* and *ALK* expression. (**C**) UMAP plots showing human fetal adrenal gland cell subpopulations as identified previously from scRNA-Seq^10^. Blue and brown color gradients indicate *ALK* and *ALKAL* expression as shown in the color key. Main cell populations labelled. (**D**) UMAP plots of the adrenocortical cluster with color gradients indicating different fetal adrenocortical signature scores as indicated. (**E**) Immunohistochemistry images of Alkal2 and pALK stained mice adrenal gland sections as indicated. Images are representative of 3 independent mice. Scale bars indicate 250 µm. (**F**) Bar plot indicating percentages of *ALKAL2* expressing primary NB tumors with high (> 90^th^ percentile), intermediate and low (<10^th^ percentile) fetal zone signatures, as indicated. RNA-seq data derived from *Cangelosi et al, 2020* ^35^. *P* values calculated using Chi-square test. (**G**) Proposed model of fetal-zone derived ALKAL2 and adrenomedullary ALK interactions putatively contributing to adrenal NB tumor formation in a subset of patients. Created in BioRender.

Our findings suggest an adrenocortical origin of these cells/spots. During fetal development, the adrenal cortex develops from the adrenal primordium (AP) and is composed of an outer definitive zone (DZ) and a larger, inner fetal zone (FZ). Postnatally, the FZ regresses and the DZ develops into 3 adult cortical zones: an outer ZG, a zona fasciculata (ZF) and an inner zona reticularis (ZR)^20,21^. To determine the precise origin of the adrenal clusters in *NB2Post* tumor samples, we compared their expression pattern with a set of fetal and adult adrenocortical gene signatures (See *Methods*)^22,23^. High similarities were observed with the fetal FZ cells and the postnatal ZF and to a lesser extent also the ZR cells, but not with the ZG cells (Fig. 2D). Based on the similarities to both prenatal and postnatal adrenocortical cells, we annotated these cells as the “Adrenocortical (AC)-like cluster”.

### Spatial expression patterns suggest paracrine interactions between ALKAL2-expressing AC-like cells and ALK-expressing neuroendocrine cells

In both sections from tumor sample *NB2Post* we observed distinct spatial regions of AC-like cells, chromaffin NE2-CA cells and NE1 cancer cells (Fig. 3A). Strikingly, the ALK ligand *ALKAL2* was one of the most upregulated genes (mean log2FC = 4.9; *P* = 3.3e-224), and, interestingly, a mutually exclusive expression pattern was observed between the 3 clusters with AC-like cells expressing *ALKAL2*, *NE1* (cancer) cells expressing its oncogenic receptor *ALK* and the NE2-CA cells expressing neither (Fig. 3B). While recent evidence from our group and others suggests a previously unanticipated therapeutically targetable driver function of this ligand in NB^24,25^, the cellular source of *ALKAL2* during early fetal development remains unclear.

To determine whether *ALKAL2* is expressed during normal adrenocortical development, we mined a previously published scRNA dataset that was obtained from the developing human adrenal gland^10^. In line with our findings, *ALKAL2* was uniquely expressed in the adrenocortical cells, while *ALK* expression was restricted to adrenomedullary cells, with the highest number of cells expressing *ALK* found in the neuroblast population (99.9%; Fig. 3C). When applying the fetal adrenocortical signatures on these data, we observed a striking overlap between *ALKAL2* expressing adrenocortical cells and cells that are predicted to have an FZ origin (Fig. 3D). Relatedly, an analysis of an independent atlas of human adrenal gland development^26^ confirmed an increasing *ALKAL2* expression (*r* = 0.63; *P* = 6.4e-03) between 40 and 75 days post conception (Fig. S5C). Interestingly a concomitant decrease in *ALK* expression was observed (*r* = −0.76; *P* = 4.1e-04).

These findings suggest that the proximity of the cortical ALKAL2-secreting cells to the ALK expressing adrenal medulla results in paracrine crosstalk during fetal and early postnatal development. To further confirm this putative cellular interaction, we performed Alkal2 and P-Alk immunohistochemical staining of postnatal 43 days old mouse adrenal glands. In line with our other findings, we detected strong Alkal2 expression in the inner layers of the adrenal cortex, (Fig. 3E). Conversely, P-Alk was uniquely expressed in the medulla. This suggests that Alk expressed in the adrenomedullary cells may be activated by Alkal2 secreted from the proximal ZR or the FZ in the postnatal or fetal adrenal gland, respectively.

To estimate the broader clinical relevance of these findings, we derived an AC-like signature from the 50 most differentially expressed genes in the AC-like cluster and applied this signature on an independent bulk RNA-Seq datasets obtained from primary NB tumors. Interestingly, 97% of the tumors with the strongest AC-like signature (>90^th^ percentile) expressed *ALKAL2*, while this dropped to 22% in tumors with intermediate signatures and 19% in tumors with the weakest signatures (*P* = 1.4e-20; Fig. 3F).

In summary, our result suggests that the interaction between adrenocortical *ALKAL2* and adrenomedullary *ALK* could play a role in the initiation and/or progression of at least a subset of adrenal NBs (Fig. 3G).

### NTRN is expressed by AC-like cells and stimulates NB cell proliferation and migration

To find support for other putative communication trajectories between AC-like cells and *NB2Post* clusters, we performed a cell-cell communication analysis. This analysis predicted 67 secreted signaling interactions that were found between the AC-like cluster and the NE1 as well as the NE2-CA cells, including the *ALKAL2* -*ALK* interaction. Additionally, interactions were predicted with several genes that were amongst the most upregulated genes in the AC-like cluster, including *FGF9* (with *FGFR1*) and *NRTN* (with *GFRA2* and *RET* ; Fig. 4A-B).

**Figure 4.**
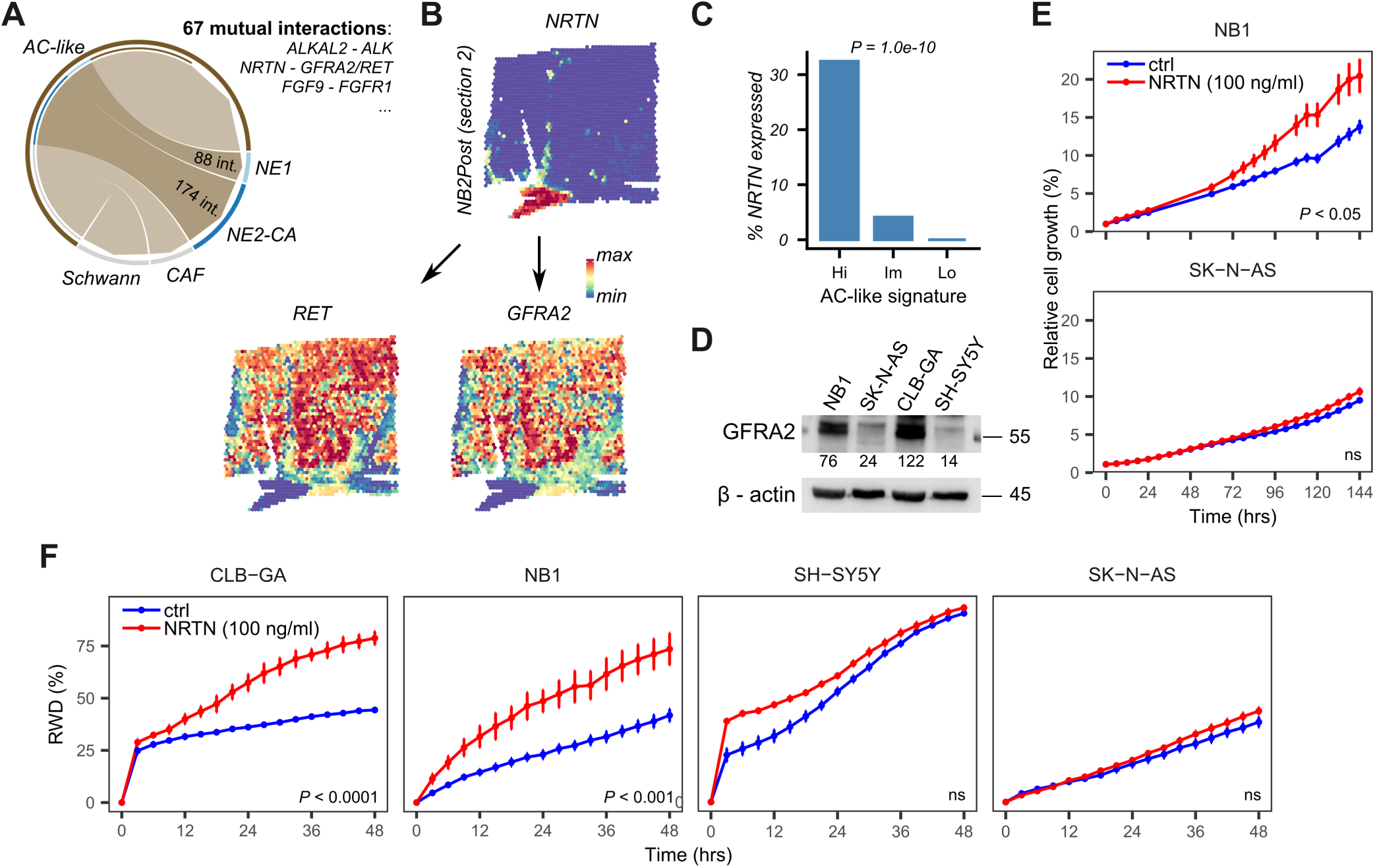
Identification of a spatially defined NRTN – GRFA2/RET interaction and experimental validation in NB cells. (**A**) Chord plot showing the number of predicted outgoing signaling interactions from the AC-like cluster. Number of interactions with NE1 and NE2-CA and examples of common interactions given. See table S3 for detailed results. (**B**) Spatial gene expression plots showing *NRTN*, *GFRA2* and *RET* expression sections as indicated. (**C**) Bar plot indicating percentages of *NRTN* expressing primary NB tumors with high (> 90^th^ percentile), intermediate and low (<10^th^ percentile) fetal zone signatures, as indicated. RNA-seq data derived from *Cangelosi et al, 2020* ^35^. *P* values calculated using Chi-square test. (**D**) Western blot showing expression of GFRA2 in four different NB cell lines, as indicated. Normalized densitometry values indicated below each blot. (**E-F**) Time-dependent effect of *NRTN* application (50 or 100 ng/ml; n = 3) on (**E**) relative wound density (RWD; measured using a scratch/wound migration assay), and, (**F**) cell growth (measured on Incucyte S3 system) on different NB cell lines as indicated. Asterisks indicate significance with *P* values calculated using an unpaired two-sided student’s t-test. *, *P* ≤ 0.05; **, *P* ≤ 0.01; ***, *P* ≤ 0.001; ****, *P* ≤ 0.0001.

*NRTN* encodes a member of the glial cell line-derived neurotrophic factor (GDNF) family of ligands that binds and activates the GFRA2-RET receptor complex^27^. While NRTN is considered an important determinant of the survival of sympathetic and sensory neurons^28^, and has been demonstrated to activate RET in NB cells previously^29^, little is known about its role in NB carcinogenesis. Interestingly, when applying the AC-like signature on independent primary NB RNA-Seq data, the percentage of NB samples that express *NRTN* was significantly higher in samples with high (32%) as compared to intermediate (4%) or low (0%) AC-like signature scores (*P* = 1.0e-10; Fig. 4C).

Based on these findings, we selected the NRTN-GFRA2/RET interaction for further experimental validation. We first confirmed the expression of the NRTN receptor GFRA2 in 4 different NB cell lines, similar to what was demonstrated for RET previously^30^. Relatively high GFRA2 expression was observed in NB1 and CLB-GA cells while SK-N-AS and SH-SY5Y cells expressed the receptor rather weakly (Fig. 4D). Stimulation of these NB cell lines with NRTN (100 ng/ml) resulted in enhanced cell migration as well as cell proliferation (Fig. 4E-F). Interestingly, the extent of this enhanced migration correlated with the relative GFRA2 expression differences between the NB cells and were more pronounced in NB1 (+ 31.75% after 48h; P = 0.0008) and CLB-GA cells (+ 34.41% after 48h; P = 0.0001) than in SK-N-AS cells (+ 5.28% after 48h; P = 0.0309). Similarly, increased proliferation was more pronounced in NB1 cells (+ 6.70% after 6d; P = 0.0086) than In SK-N-AS cells (+ 1.83% after 6d; P = 0.1926).

### Intercellular communication trajectories between niche-specific undifferentiated 11p gained neuroendocrine cells and their surrounding macrophages

In tumor sample *NB1Post* (section 2) we observed a highly specific spatial pattern with a rim of macrophages (Macro cluster) almost completely surrounding a unique and spatially distinct NE subcluster (NE4) that was not found elsewhere in the sample (Fig. 5A). Interestingly, several genes that have been previously associated with NB were amongst the 10 most upregulated genes as compared to the rest of the sample (e.g. *DUSP2*, *VGF, SCG2*) and, strikingly, 3 of these genes were located on the short arm of chromosome 11 (*WEE1*, *CALCA* and *CALCB*) (Fig. 5B-D). As there was bulk DNA sequencing data evidence for arm-level copy number gains in these chromosomes (Fig. 1A), we inferred and spatially mapped these arm-level CNVs. Both gains were confirmed and, remarkably, the 11p gain had a subclonal origin and correlated strikingly well with the NE4 clusters (Fig. 5D and Fig. S6).

**Figure 5.**
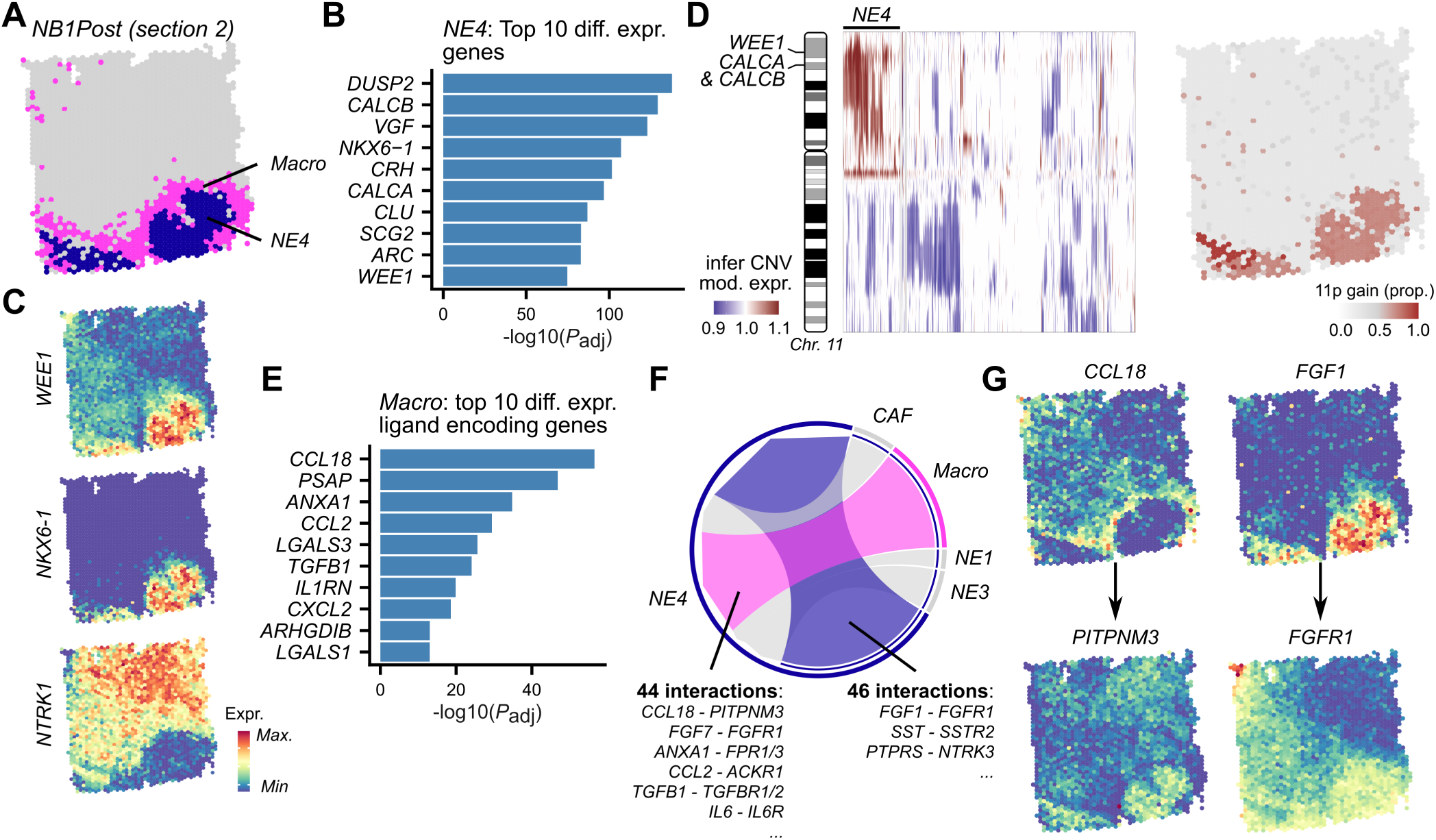
Intercellular signaling interactions between macrophages and undifferentiated, 11p amplified neuroendocrine cancer cells in *NB1Pre*. **(A)** Tumor tissue locations of the NE4 cancer cell cluster and the surrounding rim of macrophages in the second section of *NB1Pre*. (**B**) Bar plot showing the -log10(*P* _adj_) values of the 10 most upregulated genes in the NE4 cluster. (**C**) Spatial expression plots of selected genes as indicated. (**D**) Chromosome 11 copy number spot profiles, with genes that are in the top 10 upregulated genes mapped to their chromosomal location (left). Visualization of the average 11p copy number gain signal (right). Copy numbers inferred using *inferCNV*. (**E**) Bar plot showing -log10(*P* _adj_) values of the 10 most upregulated receptor ligand encoding genes in the macrophage cluster. (**F**) Chord plot showing the number of predicted intercellular incoming signaling interactions in NE4, with several examples indicated. See Table S3 for detailed results. (**G**) Spatial gene expression plots for selected ligand – receptor interaction pairs.

The 4^th^ most upregulated gene in NE4 was *NKX6-1* and the expression of this gene was confirmed to be restricted to NE4 (Fig. 5C). As this gene is generally known as a stemness marker^31,32^, and given the NE4-specific absence of expression of the bona-fide neuronal differentiation marker *NTRK1*, these results suggest that NE4 is an undifferentiated or poorly differentiated NE cluster that may have arisen in response to treatment.

To better understand the putative interactions between the rim of macrophages and the niche of NE4-cells, we next focused on the differentially expressed ligand-encoding genes in the Macro cluster. *CCL18* was the most significant upregulated gene in the macrophage cluster (Fig. 5E) and, interestingly, had a complementary expression pattern with its receptor *PITPNM3*, with the latter being mainly expressed by the NE4 cluster. A cell-cell communication analysis was performed to find additional evidence for ligand-receptor signaling interactions between macrophages and NE4 cells. Forty-four significant (*P* < 0.05) intercellular communication trajectories were identified between the Macro and the NE4 cluster, including between the *CCL18* ligand and *PITPNM* receptor and several other cytokine receptor pairs (*CCL2* – *ACKR1* and *IL6* - *IL6R* ; Fig. 5F-G and Table. S3). We also identified several interactions with the NE4-expressed oncogene receptor *FGFR1*, both with ligands predicted to be produced by the macrophages (*FGF7*) and with ligands produced by NE4 cells (*FGF1/2*) (Fig. 5G).

### CCL18 enhances PITPNM expressing NB cell migration

While *CCL18* - *PITPNM3* interactions have been previously demonstrated to promote invasion, metastasis and chemoresistance in breast cancer^33,34^, nothing is currently known about their role in NB. Therefore, we experimentally assessed the phenotypic effects of CCL18 in NB cell lines. We first confirmed the expression of the PITPNM3 protein using western blotting in 4 different NB cell lines. Higher expression was found in NB1 and CLB-GA cell lines as compared to SK-N-AS and SH-SY5Y (Fig. 6A). Given the previously suggested role of CCL18 on invasion and metastases, we explored the effects of this chemokine on NB cell migration in NB1 and SK-N-AS cells. Interestingly, CCL18 (100 ng/ml) significantly enhanced migration in both cell lines (*P* < 0.05). Additionally, we also observed a moderately increased cell proliferation in NB1 cells but not in SK-N-AS cells (Fig. 6B-C).

**Figure 6.**
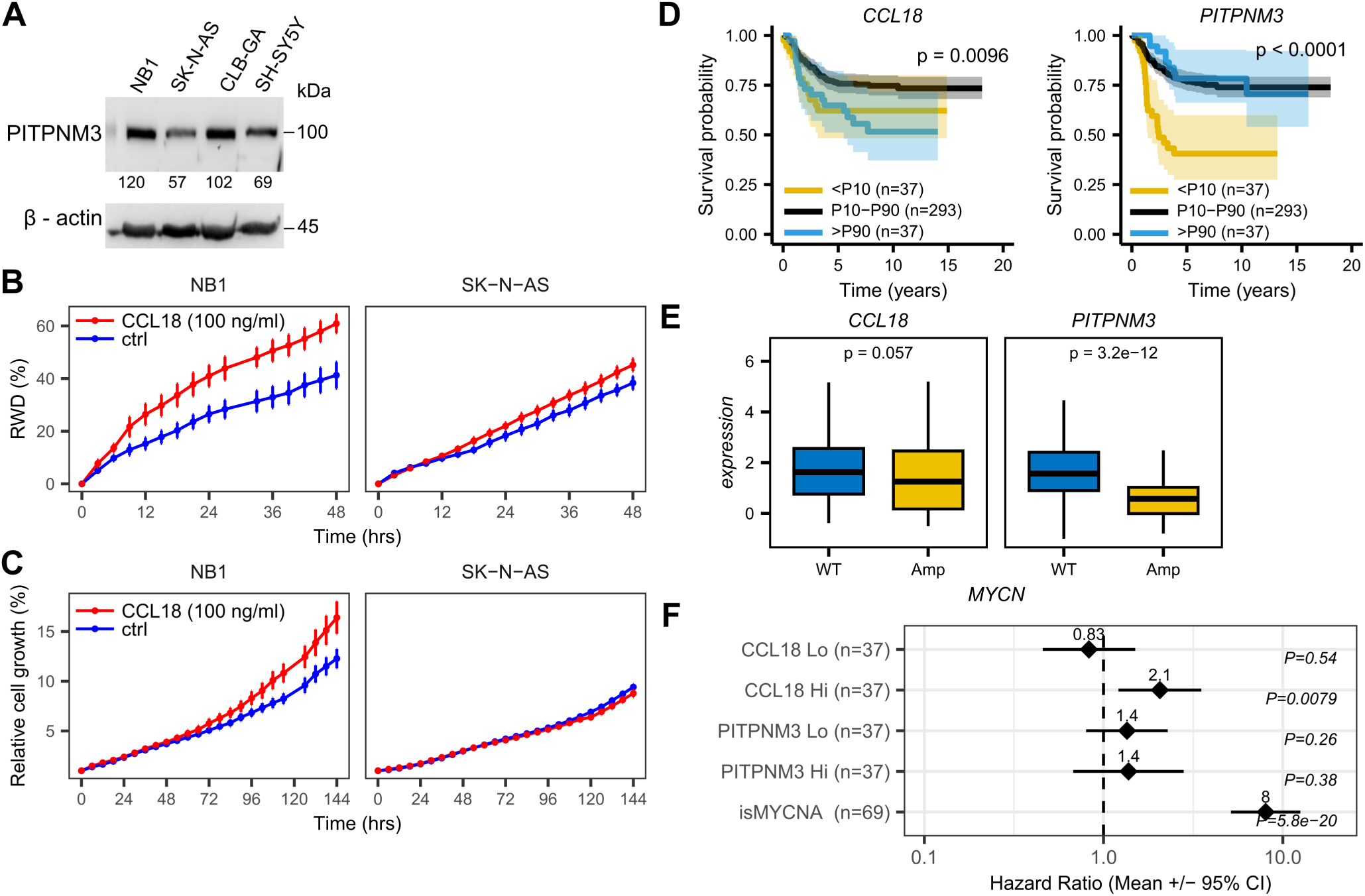
Experimental and clinical validation of the influence of CCL18 on NB cells. (**A**) Western blot showing the protein expression profile of the CCL18 receptor PITPNM3 in 4 NB cell lines as indicated. Normalized densitometry values indicated below each blot. (**B-C**) Time-dependent effect of *CCL18* application (100 ng/ml; n = 3) on (**B**) relative wound density (RWD; measured using a scratch/wound migration assay), and, (**C**) cell growth (measured on Incucyte S3 system) in different NB cell lines as indicated. Asterisks indicate significance with *P* values calculated using an unpaired two-sided student’s t-test. *, *P* ≤ 0.05; **, *P* ≤ 0.01; ***, *P* ≤ 0.001; ****, *P* ≤ 0.0001. (**D-F**) Survival analysis of CCL18 and PITPNM expression in 364 primary NB RNA-Seq data^35^. (D) Kaplan-Meier survival plots comparing overall survival between patients with high (>90^th^ percentile, intermediate (10^th^ – 90^th^ percentile) and low (<10^th^ percentile) *CCL18* and *PITPNM3* expression as indicated. *P* value calculated using log rank test. (**E**) Boxplots comparing *CCL18* (left) and *PITPNM3* (bottom) expression between *MYCN* amplified and *MYCN* wild-type tumors. *P* value calculated using two-sided unpaired Wilcoxon’s test. (**F**) Forest plots comparing hazard ratios +/− 95% confidence intervals for 5 variables as indicated. Results were obtained using a Cox proportional hazards multivariate regression analysis

To determine whether these findings have potential clinical relevance, we focused on a large dataset of transcriptomic and associated clinical NB data^35^ (Fig. 6D-F). Interestingly, the expression of *PITPNM3* was strongly reduced (*P* = 3.2e-12) in NB patients with MYCN amplifications, one of the main genomic determinants of high-risk NB (Fig. 6E). A similar trend was observed for *CCL18*, although this was not significant (*P* = 0.057). To determine the prognostic value of these genes, we performed a multivariate Cox logistic regression analysis and associated *CCL18* and *PITPNM3* expression as well as MYCN amplification status to observed survival. Interestingly, high expression of CCL18 (i.e., > 90^th^ percentile) was significantly associated with worse survival (HR = 2.1 as compared to patients with intermediate expression, i.e., 10^th^ – 90^th^ percentile; P = 0.0079), independent of *MYCN* amplification status (HR = 8.0 for patients with *MYCN* amplification as compared to *MYCN* wild-type; P = 5.8e-20: Fig. 6F).

## DISCUSSION

Neuroblastoma is the most frequent extra-cranial tumor in the pediatric population and accounts for 15% of pediatric cancer-related death. The tumor exhibits a large degree of ITH which is one of the main causes of treatment failure and relapse. While valuable insights in ITH have been obtained from sequencing of multifocally sampled tumor regions or from scRNA-Seq studies, these studies generally lack the full spatial tissue context, restricting the conclusions about neighboring intercellular interactions and paracrine communication trajectories. Here we addressed this limitation by applying the Visium spatial transcriptomics platform on 6 sections from 3 tumor samples obtained from 2 NB patients. Despite our study’s relatively low sample size, we demonstrated that important insights can be gained into subclonal copy number variations (CNVs) and into novel paracrine interactions between unique spatial niches of cells in the NB tumor landscape.

We first clustered and annotated the spatial transcriptomics data. This annotation was challenging due to the oligocellular spatial resolution of the applied Visium platform. Transcriptomic signals were derived for circular tissue spots with a diameter of 55 µm, implying a transcriptomic mix of 1-10 cells. Therefore, we annotated the regions based on the dominant signal that was derived using several criteria: differential gene expression of marker genes, gene set enrichment analysis, similarities to previously annotated single cell data and copy number variability (which is mainly expected in cancer cells). For most clusters the evidence clearly identified one dominant cell type based on all criteria, while other clusters were more mixed. This was illustrated by the NE1 cluster in the *NB1Post* sample whose annotation was solely based on the high copy number variability but is likely strongly mixed with other cell types.

The cancer cell population in NB tumors exists in two main cell states, adrenergic or mesenchymal^14,15^. Tumor cells in the mesenchymal state are more chemo-resistant and are believed to be the main drivers of post-therapy relapse^15,36^. However, the spatial organization of these cell states are underexplored in NB. Therefore, we developed a spatially applicable, single score based on the adrenergic and mesenchymal gene signatures and visualized the complete cellular state spectrum in our 6 tumor sections. While the majority of the NE cancer cell clusters were in a more adrenergic state, co-existence with NE cells in more mesenchymal states was confirmed in each tumor. This variation in cell state proportions aligns with recent scRNA studies, which show adrenergic NE populations as the dominant group compared to mesenchymal state^37,38^. Furthermore, the cells/spots were found along a full continuum of adrenergic to mesenchymal transcriptional states, suggesting that cell states represent a differentiation spectrum rather than discrete entities. This highlights the complexity and plasticity of the transcriptional ITH in NB, where cells may span continuums of transcriptional programs, reflecting a range of cell states. One critical remark here is that the cell state signal could be influenced by non-cancer cells, both from within the same spots (mixtures of cells) and from nearby spots due to diffusion of mRNA (spot swapping). It is currently unclear how this could have affected our analysis and future analyses on higher resolution spatial transcriptomics platforms will be required to confirm this adrenergic to mesenchymal differentiation spectrum.

In the first patient’s chemotherapy-treated tumor (*NB1Post*), we identified a unique spatial niche of undifferentiated cancer cells that was completely isolated from the rest of the sample by a peripheral rim of macrophages. Strikingly, these cells harbored a cluster-specific copy number gain in the short arm of chromosome 11 (11p) with several genes in the top differentially expressed genes, including *WEE1*. *WEE1* encodes a dual-specificity kinase that inhibits the activation of CDK1 and CDK2, thereby regulating cell cycle progression and DNA damage responses^39,40^. These findings suggest that this 11p gain, likely in combination with other subclonal CNVs (e.g., we also identified an 8q gain) provide a selective advantage in this tumor cluster, resulting in chemotherapy resistance. A recent study identified 11p/proximal 11q gains in a subgroup of chemotherapy-treated MYCN-non-amplified NB tumors using single nucleus RNA-Seq and whole-genome sequencing^37^. This copy number alteration is strikingly similar to that observed here in *NB1Post* and, remarkably, was similarly associated with increased stemness. We also identified *CCL18* as the top differentially expressed receptor ligand in the peripheral macrophage population. A cell-cell communication analysis identified the interaction of *CCL18* with the receptor *PITPNM3* that was expressed by the undifferentiated cancer cell cluster. We experimentally validated the stimulatory effect of CCL18 on NB cellular migration, possibly in line with the invasion and metastasis properties that have been described for the CCL18-PITPNM3 interaction in breast cancer^33,34^. Interestingly, using independent clinical NB data we demonstrated an adverse effect of high *CCL18* expression on NB prognosis (Hazard Ratio = 2.1), independent of *MYCN* amplification status. Additionally, the identification of a potential FGF1/FGFR1 signaling axis in this unique undifferentiated cancer cell cluster is of particular interest, as a recent report identified a role for FGFR signaling in lorlatinib resistance in NB^41^, offering therapeutic opportunities.

One of the key findings of our study was the identification of an adrenocortical stromal component of the NB TME, which we termed AC-like cells. Adrenocortical cells have been identified in single cell studies previously but have been largely ignored based on the assumption they were adrenal contaminants^37,38,42^. The unique spatial pattern of AC-like nests and in stripes that were scattered throughout the tumor, strongly argue against an incidental adrenal contamination. Additionally, the main AC-like zone cluster was located adjacent to a clearly demarcated *PNMT* expressing adrenomedullary chromaffin cell cluster, which was fully integrated in the tumor, suggesting a complex interplay between AC-like cells, CA cells and NE cancer cells. We identified several paracrine interactions between these different regions. The most interesting was between *ALKAL2* and *ALK* that were uniquely expressed in the AC-like cells and the NE cancer cells, respectively. ALK signaling potentiates *Th-MYCN* driven NB development^24,43^, and its absence significantly reduces tumor penetrance in *Th-MYCN* driven mouse models of NB^44^. Together our data raise the intriguing possibility that NB tumors co-opt ligand producing AC-like cells to maintain tumorigenic potential.

Our results provide putative novel insights into early adrenal NB carcinogenesis, where *ALKAL2* secreting adrenocortical fetal zone cells could stimulate *ALK* activity in the adrenal medulla, supporting the initiation and progression of NB (Fig. 3G). Apart from the evidence we provide in our study, this model could also explain why non-adrenal, thoracic NB tumors are more likely to have gain-of-function *ALK* mutations than adrenal NB^4^. Indeed, if adrenal NB is driven by ALKAL2 in some circumstances, the selective pressure on *ALK* mutations would be lower than in the absence of ALKAL2. If our model is correct, one of the key questions would be which permissive conditions would be required for ALKAL2 to initiate NB formation.

In conclusion our whole transcriptome ST approach confirmed the strong transcriptional and cellular intratumoral heterogeneity of NB. We identified several unique spatial niches and predicted paracrine interactions between these niches. *CCL18* expressing macrophages completely surrounded a tumor-specific 11p/proximal 11q amplified, undifferentiated and *PITPNM3* expressing cancer cell cluster. Lastly we propose a role for ALKAL2 expressing fetal zone cells in adrenal NB carcinogenesis during development. Overall, this study provides novel insight into the highly dynamic nature of the NB tumor microenvironment and identifies multiple signaling axes with therapeutic potential for future exploration.

## METHODS

### Patients and tumor samples

Formalin fixed paraffin-embedded (FFPE) tumor samples were obtained from the Section of Pathology, Department of Clinical Genetics, Pathology and Molecular Diagnostics, Office of Medical Services, Lund, Sweden, after informed consent for research under the ethics approval 2023-01550-01 from the Swedish Ethical Review Authority. Patient selection was done by tracing an ongoing research cohort of NB patients back in time from 2023 and selecting the first patients with tumor material of sufficient quality and quantity for the analyses.

Two 5 µM tumor sections were obtained from 3 FFPE embedded tumors from 2 high-risk NB patients (*NB1* and *NB2*): a treatment-naïve (*NB1Pre*) as well as a chemotherapy treated (*NB1Post*) *MYCN* wild type tumor and a chemotherapy treated *MYCN* amplified tumor (*NB2Post* ; Fig. 1A). The 6 tumor sections were subjected to spatial transcriptomic (ST) sequencing using Visium Spatial Gene Expression FFPE assay, following the instructions from the manufacturer (10X Genomics). *NB1* tumor sections were directly placed onto Visium v1 slides for probe hybridization. For *NB2* tumor samples, probe hybridization and ligation was first performed on standard frosted glass slides, followed by transfer of the hybridized probes to Visium v2 slides using the CytAssist instrument. Gene expression libraries were prepared according to the manufacture’s protocol and sequenced on NextSeq2000. The sequenced reads were demultiplexed and mapped to the reference genome, GRCh38 (build 2020-A, 10X Genomics), using *Space Ranger* software v1.3.1 (10X Genomics).

### Spatial transcriptomics data processing

ST data was processed and analyzed using *R* v4.2.2 and the *R* package *Seurat* v5.0.0.^45^ Firstly, the *Space Ranger* output files (raw expression matrices and spatial metadata) were loaded using the functions *Read10XRaw* and *read10xSlide* from the R package *SpotClean* v1.5.2^46^. The *SpotClean* algorithm was used to adjust for spot swapping, a phenomenon where unique molecular identifiers (UMIs) bleed from nearby spots and contaminate neighboring spots. The resulting data was converted to Seurat objects for further processing.

Individual samples were normalized via Seurat’s *SCTransform* workflow, followed by merging of the two replicate sections from the same tumor (i.e., *NB1Pre*, *NB2Pre*, *NB2Post*). Unsupervised Louvain clustering was subsequently performed via *FindCluster*s.

Cell population annotations were based on hematoxylin and eosin staining (HE) sections, multiple clustering resolution testing and visualization with the *clustree* v0.5.1^47^ package and the top differentially expressed genes in each cluster. CNV calling was performed with *inferCNV* v1.20.0, using raw counts as input and all spots not belonging to NE clusters as reference group. The cutoff parameter was set to 0.1. Spot identities were further investigated through integration with NB scRNA-Seq atlas (NBAtlas)^18^ or fetal adrenal gland scRNA-Seq^10^ data, using either Seurat’s label transfer or *ClusterFoldSimilarity* (v0.99.14) approach^49^. *SpatialDimPlot* and *SpatialFeaturePlot* functions were used to visualize the cell clusters and gene expression level in the ST data.

### Imputation of missing values/technical zeroes

The gene expression matrix of *NB1* tumor samples (*NB1Pre*, *NB1Post*), processed with Visium v1, contained large number of technical zeros. To improve visualization of spatial features, we imputed missing values using the *ALRA* R package^50^.

### ST and scRNA-Seq signature score and similarity analyses

Gene signatures were obtained from publicly available datasets. Adrenergic and mesenchymal gene signatures were obtained from the study by van Groningen and colleagues^15^. Gene markers for the fetal and postnatal adrenocortical zones were obtained from the top 50 differentially expressed genes as reported in 2 previous studies^22,51^.

Gene signatures were scored using the *AddModuleScore_UCell* function from the *UCell* v2.6.2 R package^52^. *ClusterFoldSimilarity* R package was used to test the similarity between clusters of different datasets^49^.

The adrenergic and mesenchymal UCell scores were normalized to their mean value within each tumor. A single spot/cell state visualization score was calculated by taking the log2 ratio of the normalized adrenergic score over the normalized mesenchymal score. This way spots with positive and negative values represent relatively higher adrenergic and mesenchymal states, respectively.

### Differential gene expression and ranked gene set enrichment analyses (GSEA)

Wilcoxon rank-sum test-based differential gene expression (DGE) analysis between respective ST spot clusters and all other spots was performed using *FindAllMarkers* function of the *Seurat* R package. Gene lists were ranked based on DGE *P* value. Reactome and PanglaoDB gene sets were downloaded from *MSigDB* v2023.1^53^ and *PanglaoDB* ^54^, respectively. GSEA was performed using the *fgsea R* package via the package *clusterProfiler* v4.6.4^55^.

### Cell-cell communication analysis

Cell-cell communication analysis was performed using the *CellChat* (v2.1.1) R package^56,57^. In brief, individual CellChat objects were created from each merged Seurat object (cf. ST data processing), using the SCT-normalized count gene expression matrix, spatial coordinates of spots and metadata. The CellChat human ligand-receptor databas included in the package was used. Ligand-receptor pairs that were identified in this study and were missing from the CellChat database (i.e., ALKAL1/2 – ALK and CCL18 - PITPNM3) were appended. Only ligand-receptor interaction pairs with annotation ‘Secreted Signaling’ were used in the analysis. Cell-cell communication networks were inferred using the truncatedMean method, with parameters ‘trim’ at 0.2, ‘interaction.range’ at 250 µm, contact.dependent = FALSE for the *ComputeCommunProb* function. Communications involving less than 10 spots were filtered out.

### Bulk RNA-seq and survival analysis

Human NB RNA-seq (batch corrected log normalized counts) and related clinical data were obtained from the study of *Cangelosi et al.* (Cangelosi et al., 2020) (available at https://www.ncbi.nlm.nih.gov/pmc/articles/PMC7563184/bin/cancers-12-02343-s001.zip). An AC-like gene signature was derived from the top 50 DEGs in the AC-like cluster (Table S3). An AC-like signature score was then calculated for each patient using the *ssGSEA* function of the *singscore* v1.22.0 R package. Patients were stratified in low, intermediate and high AC-like groups based on 10^th^ and 90^th^ percentile cut-offs and the proportion of tumor samples that expressed *ALKAL2* or *NRTN* expression (i.e., log normalized counts >1) was determined for each group. Statistical differences between the groups were assessed using a Chi-squared test.

### Cell culture, proliferation and migration assays

Cell lines used were CLB-GA, NB1, SK-N-AS and SHSY5Y. Cell lines were cultured in complete media, RPMI 1640 supplemented with 10% fetal bovine serum (FBS) and a mixture of 1% penicillin/streptomycin at 37⍰°C and 5% CO_2_.

For proliferation and growth analysis, cells were seeded, at cell density between 3000 cells/well of 96-well plate. Cells were then cultured overnight, followed by treatment with with 100 ng/ml of CCL18 (also known as MIP-4; GIBCO, Cat# PHC1254,) or NRTN (PeproTech, Cat# 4501120UG,), depending on the cell line. The growth mediums, RPMI 1640, was supplemented with 1% FBS during cell treatment. Cell confluency/proliferation was monitored live using the Incucyte S3 Live Cell Analysis system (Essen BioScience) for several days as indicated. Rate of cell growth under all conditions were determined using the Incucyte S3 software.

For cell migration analysis, 70,000 cells were seeded per well, into the Incucyte Imagelock 96-well Plate from Satorius (Germany). Cells were then cultured overnight. Scratch wounds were made using the Incucyte Woundmaker tool. Cells were then gently rinsed with 1X PBS solution, followed by addition of RPMI1640 growth medium supplemented with 1% FBS, with or without CCL18 or NRTN at the indicated concentrations. Cell migration was monitored live using the scratch wound module of the Incucyte S3 Live Cell Analysis system. Relative wound density was determined and visualized suing the Incucyte S3 software.

### Immunoblotting analyses

Cells were lysed in 1X Laemmli sample buffer and heated for 10 minutes at 95°C. Samples were subjected to western blot analyses using the following antibodies PITPNM3 (1:1000) (Thermo Fisher Scientific Cat# PA5-21903, RRID:AB_11154802), GFRA2 (1:1000) (Proteintech Cat# 21973-1-AP, RRID:AB_11124728) and beta-actin (1:10,000) (Cell Signaling Technology Cat# 4970, RRID:AB_2223172). Chemiluminescence detection was done using Odyssey Fc Imager (LI-COR). Immunoblots were quantified using Image Studio Lite (v5.2) software.

### Immunohistochemistry of mouse adrenal glands

Adrenal glands were collected from 3-4 male mice of an age of 43 days on a 129X1/SvJ background. Adrenal glands were fixated in 10% neutral buffered formalin and subsequently mounted in paraffin blocks by standard histology methods. All animal experiments were completed in agreement with the Regional Animal Ethics Committee approval, Jordbruksverket (Dnr 5.8.18-02319/2023).

Antigen retrieval with a citrate buffer (pH 6.00) were performed on 4 μm sections. The sections were blocked with Blocking reagent (Roche, 11096176001) and stained with either anti-pAlk Y1507 (Abcam, ab73996, 1:400) or anti-Alkal2 (In house antibody, 1:800). An HRP-labeled anti rabbit antibody was used as secondary reagent (Jacksson, 711-035-152) and the immunoreactions were visualized using Liquid DAB+ substrate chromogen system (Dako K3468). Nuclei were counter stained with hematoxylin. Images of the slides were obtained by scanning the slides in NanoZoomer-SQ (Hamamatsu). The images were cropped in Affinity designer 2 (2.5.5).

### Data and code availability

Seurat data objects will be deposited in Zenodo. Source code used to generate the manuscript results is available at GitHub https://github.com/CCGGlab/ST_NB.

## ACKNOWLEDGMENTS

This project was funded by grants from the Swedish Childhood Cancer Foundation (TJ2021-0068 - JTS; PR2022-0029 - RHP), the Assar Gabrielsson’s foundation (FB23-92-JTS), the Swedish Cancer Society (CAN21/1459 - RHP), the Research Foundation Flanders (FWO; FWO.OPR.2023.0063 – JvdE and FWO.3F0.2024.0075 - PM), Belgian Foundation Against Cancer (STI.STK.2023.0017 – JvdE), the Swedish Cancer Society (21 1383 Pj - DG), the Swedish Childhood Cancer Foundation (PR2022-0026-DG), and Regional Clinical Research grants (ALF Projekt0067 -DG). The authors acknowledge support from the National Genomics Infrastructure in Stockholm funded by Science for Life Laboratory, the Knut and Alice Wallenberg Foundation and the Swedish Research Council, and SNIC/Uppsala Multidisciplinary Center for Advanced Computational Science for assistance with massively parallel sequencing and data delivery via the UPPMAX computational infrastructure.

## AUTHOR CONTRIBUTIONS

JTS, DGN, RHP, and JVdE conceived and designed spatial transcriptomics experiments. JTS, MB, CJ and JK prepared FFPE tissue slides and libraries for sequencing. PM performed InferCNV copy number analyses. JTS and JVdE performed the other bioinformatic analysis. JTS performed the in-vitro cell culture assays. MB, DEL and EJ performed immunohistochemistry experiments. JTS and JVdE wrote the manuscript, followed by inputs from all other authors.

## SUPPLEMENTARY FIGURE LEGENDS

**Figure S1.**
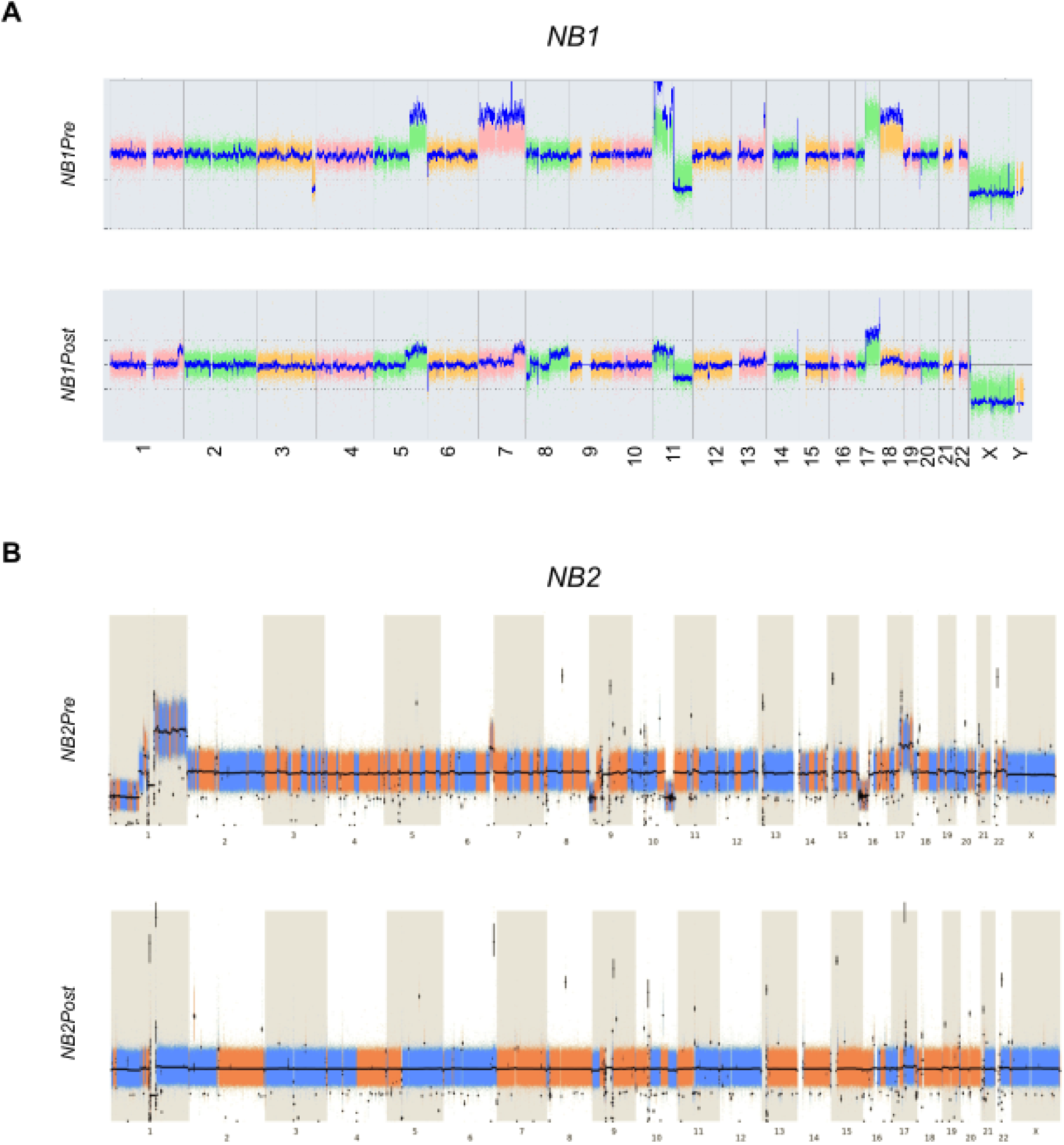
Copy number profiles from the analyzed NB tumors. Copy number profiles from NB1 (A) and NB2 (B) tumors before and after chemotherapy, as indicated. X axis labels indicate chromosome numbers.

**Figure S2.**
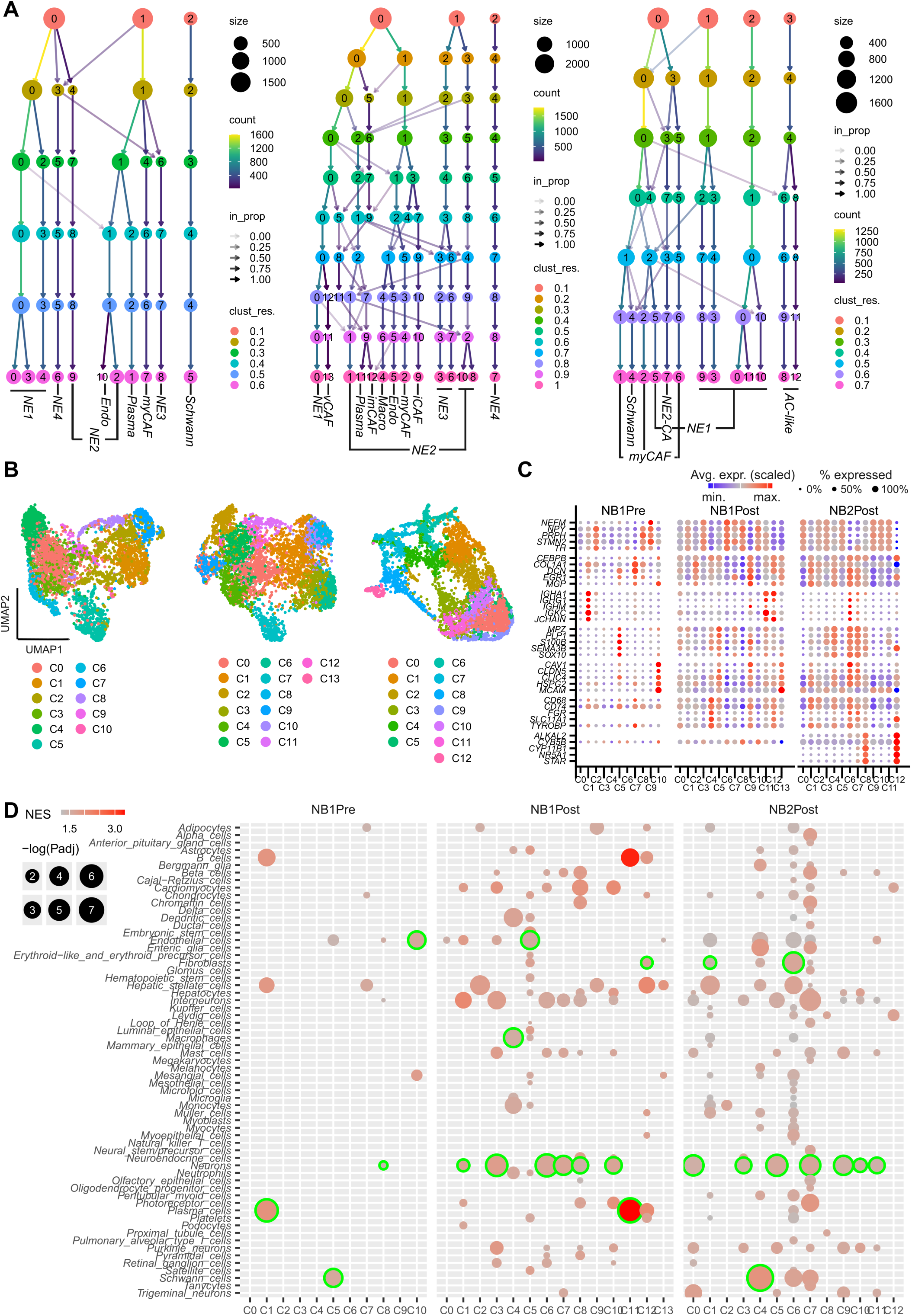
Unsupervised clustering and annotation of the 3 NB tumor samples. (**A**) Cluster tree showing the different Louvain clusters as a function of the cluster resolution (color key). The last leaf represents the resolution selected for further analysis. Cluster annotations labelled to each branch. (**B**) UMAP plot with indication of the Louvain clusters. (**C**) Dot plots showing relative expression (colors) and proportional spot expression (sizes) for the 5 genes as in main Fig. 1D. (**D**) Dot plot showing the results of a GSEA on all Louvain clusters (x axis labels) using cell-specific gene sets obtained from *PanglaoDB* (y axis labels). Label sizes and colors correspond to -log(*P* _adj_) and normalized enrichment scores (NES), respectively (see color key on top left). Green circles indicated the cell types that were used for the main cluster annotations.

**Figure S3.**
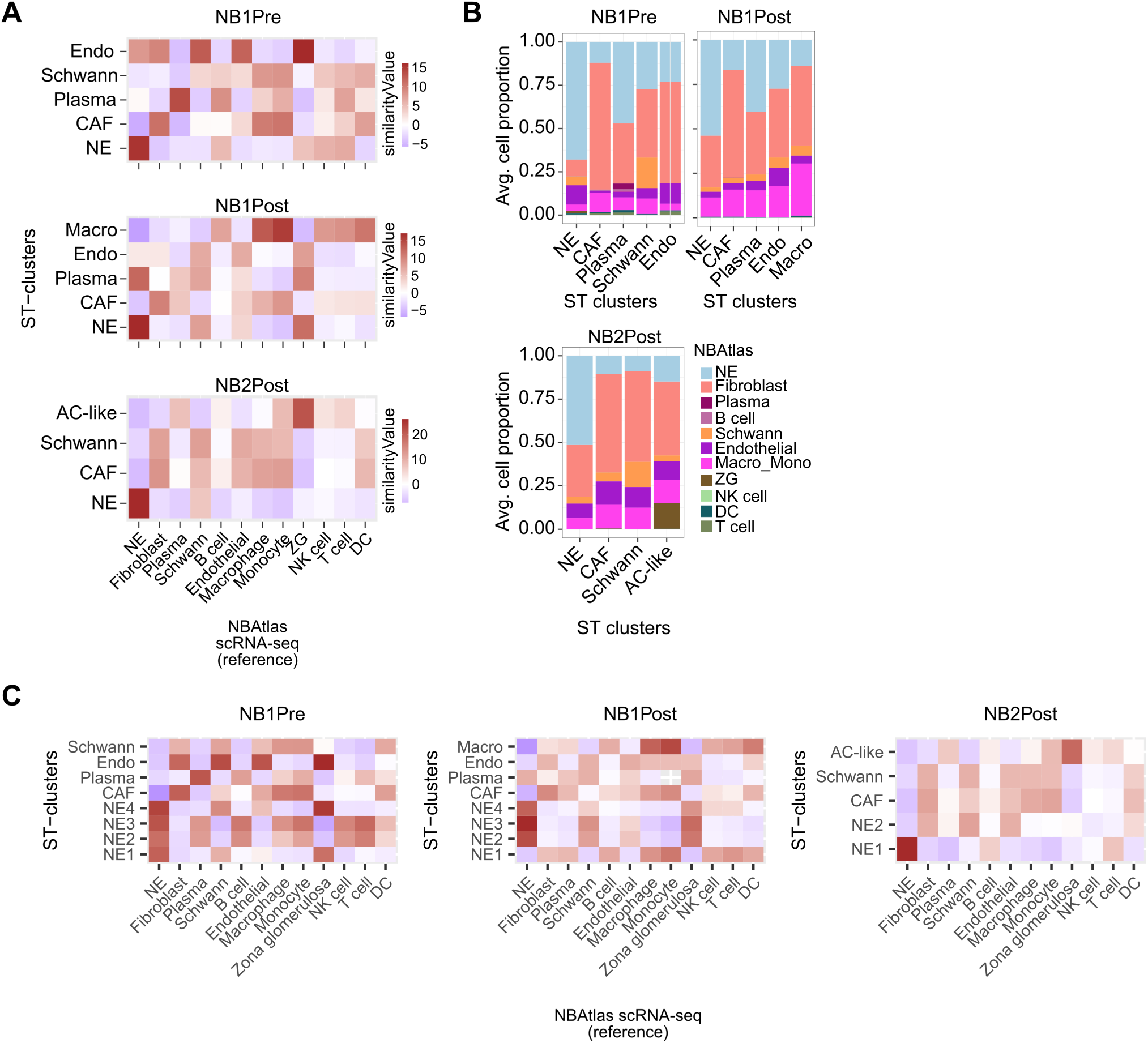
Cellular deconvolution results of the 3 NB tumor samples. Deconvolution was performed using the robust cell-type decomposition (RCTD) method and scRNA reference data obtained from the NBAtlas. (**A, C**) Heatmaps showing the relative similarities between the scRNA annotation (x axis) and the main (**A**) or NE subclusters (**C** ; y axis). (**B**) Stacked bar plots showing RCTD estimated average cell proportion in the main clusters as indicated in x axis.

**Figure S4.**
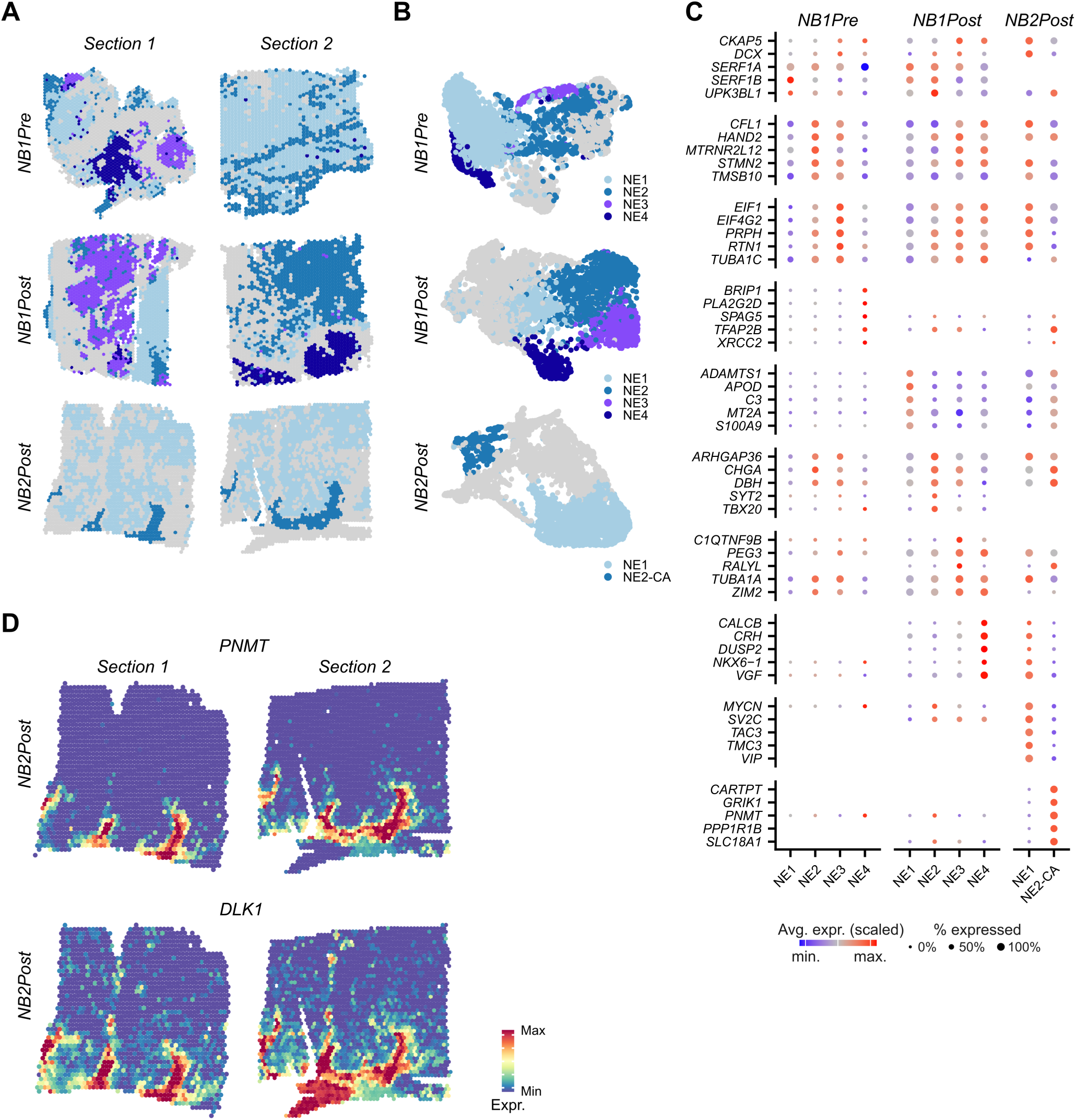
Gene expression profiles of NE subclones. The NE cluster of each tumor was subdivided in 2-4 tumor-specific subclusters (NE1 - NE4) based on spatial location and expression profile similarities. (**A-B**) Spatial location (**A**) and UMAP plots (**B**) of the different subclusters as indicated. Grey spots indicate non-NE annotated spots. (**C**) Dot plots showing relative expression (dot color) and proportional expression (dot size) of the 5 most differentially expressed genes in the NE annotated subclusters. (**D**) Spatial feature plots showing gene expression of the adrenomedullar marker *PNMT* and the adrenal marker *DLK1*.

**Figure S5.**
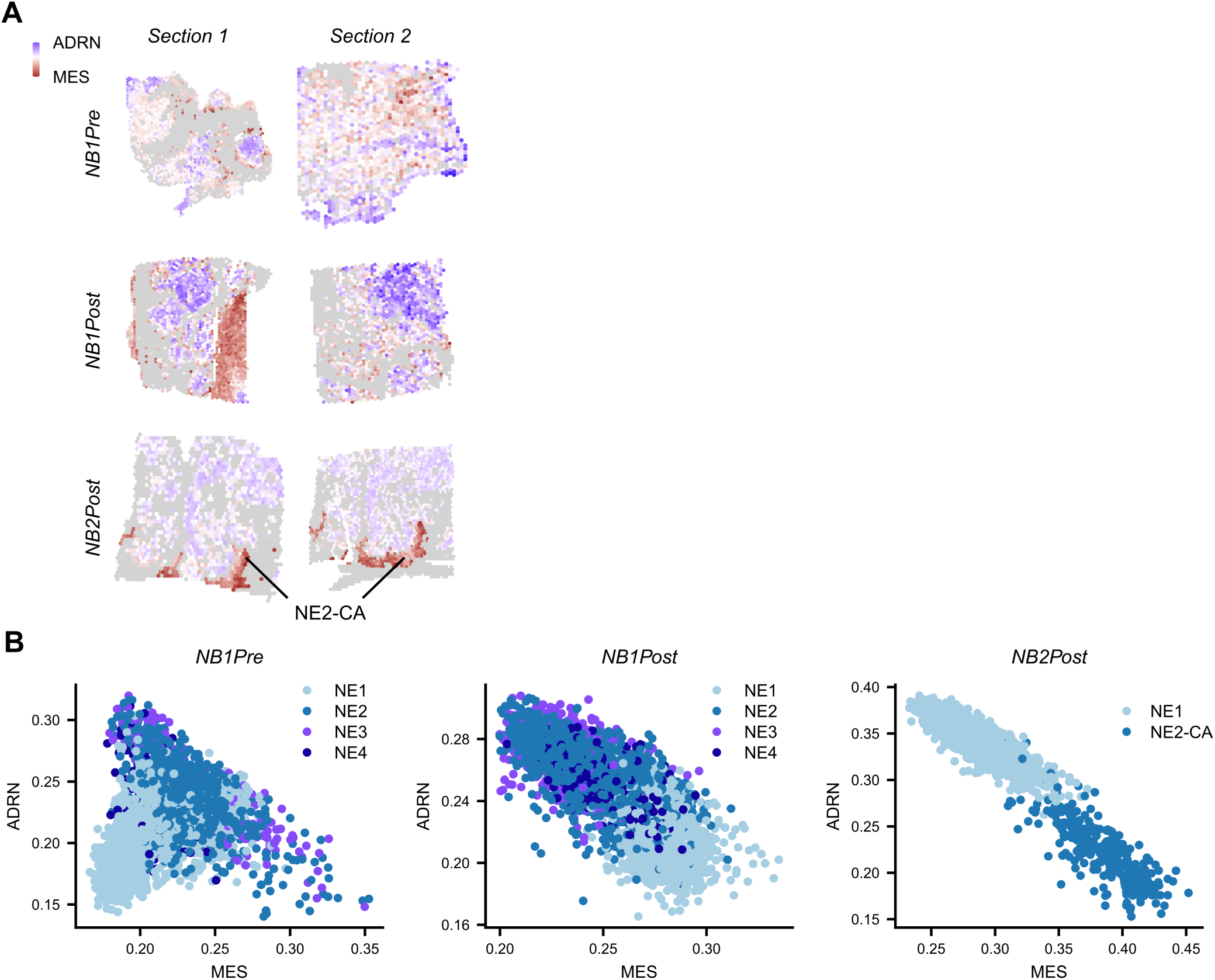
Mesenchymal and adrenergic states of the NE subclones. (**A**) Spatial plots showing the relative adrenergic or mesenchymal cell state for all samples, as indicated by the color key. (**B**) Scatter plots showing adrenergic and mesenchymal scores for each spot. Dots colored by subclone as indicated in color legend.

**Figure S6.**
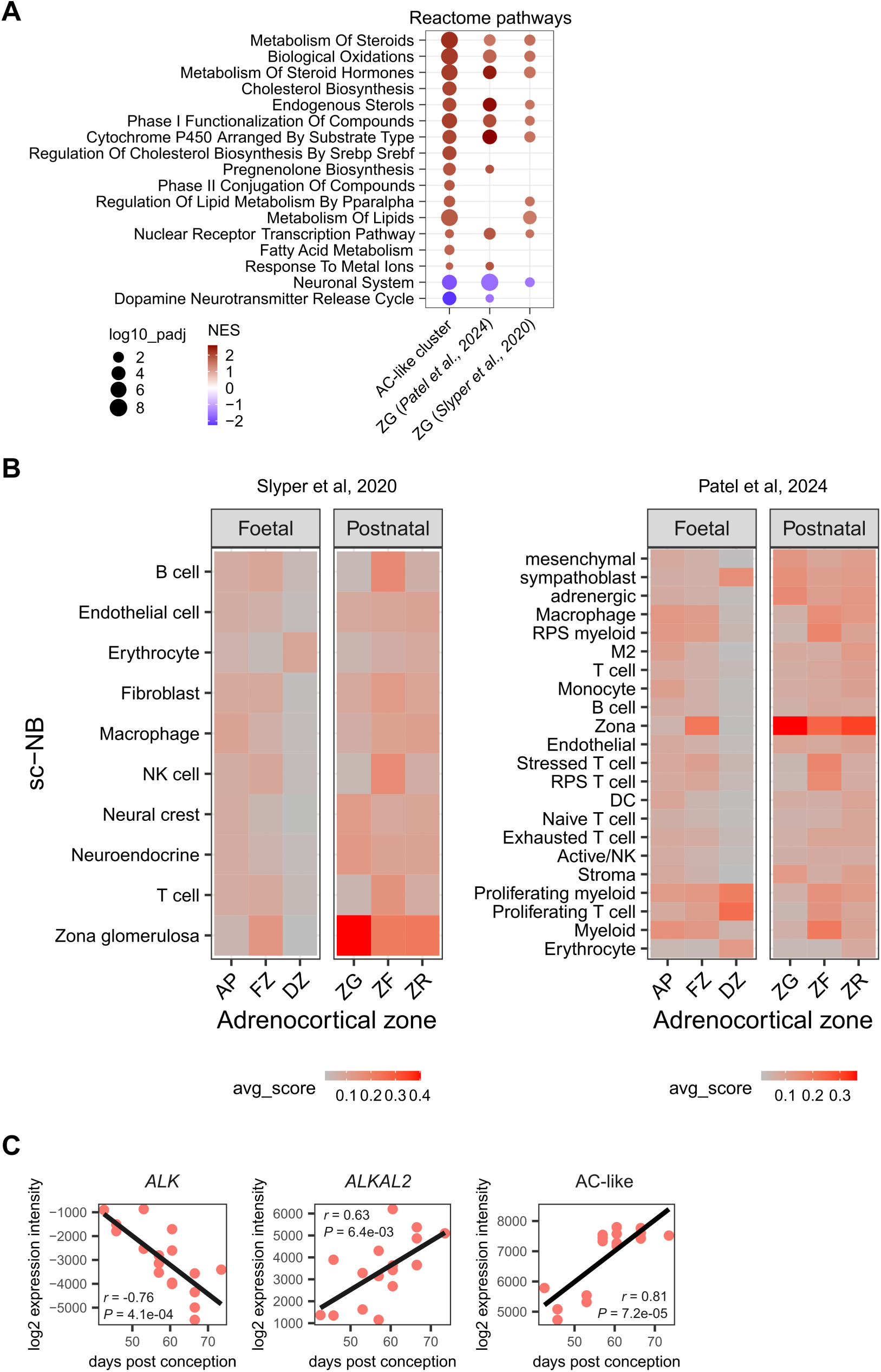
Adrenocortical signature analysis of independent human transcriptomics datasets. Adrenocortical signatures were analyzed in independent human transcriptomics studies^38,42^ (**A**) Dot plot comparing GSEA results of selected Reactome gene sets in our study with 2 scRNA-Seq studies. Dot sizes and colors correspond to normalized enrichment scores (NES) and *P* values, as indicated by color key. See table S2 for complete GSEA results. (**B**) Heatmaps showing UCell scores of fetal and postnatal adrenocortical cell type signatures on the clusters that were described by both studies. AP, adrenal primordium; FZ, fetal zone; DZ, definitive zone; ZG, zona glomerulosa; ZF, zona fasciculata; ZR, zona reticularis. (**C**) Scatter plots showing the correlation between expression of *ALK*, *ALKAL2* and the AC-like expression signature as function of time during human adrenal gland development. Linear regression line and Pearson’s correlation coefficient and *P* value indicated. Data derived from *Del Valle et al., 2022* ^23^.

**Figure S7.**
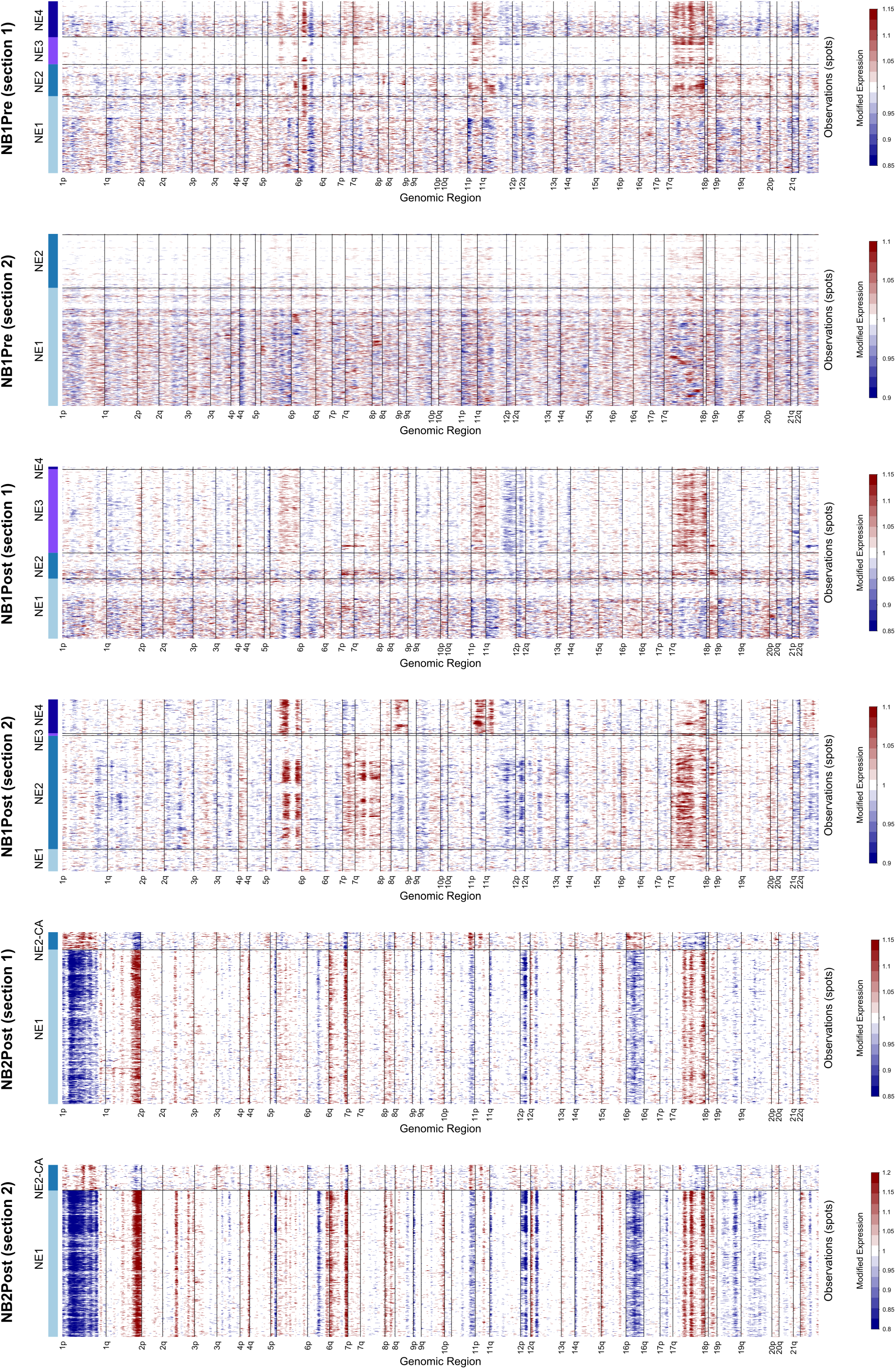
Spatial transcriptomics-based CNV profiles of the 6 NB samples. Full copy number profiles as a function of chromosome location (indicated in x axis) for all NE clusters (y axis) of the 6 analyzed tumor samples as indicated. Non-NE annotated spots were used as a reference. Copy numbers inferred using *inferCNV*. Legend key shown on the right of each plot (red = copy number gain; blue = deletion).

